# A Chemostat-Based Model for Growing Bacterial Biofilms

**DOI:** 10.1101/2025.06.22.660958

**Authors:** Fabrizio Spagnolo, Iñigo Caballero, Alexandra Goldblatt, Michael J. Loccisano, Mazharul I. Mahe, Yaxkyn Mejia, Naya Melvani, Aliza Nagel, Aiden Stanciu, Sherin Kannoly, John J. Dennehy, Monica Trujillo

## Abstract

Biofilms are groups of microbes that live together in dense communities, often attached to a surface. They play an outsized role in all aspects of microbial life, from chronic infections to biofouling to dental decay. In recent decades, appreciation for the diversity of roles that biofilms play in the environment has grown. Yet, most bacterial studies still rely upon approaches developed in the 19^th^ century and center on planktonic populations alone. Here we present a chemostat-based experimental platform to investigate not only biofilms themselves, but how they interact with their surrounding environments. Our results show that biofilms grow to larger sizes in chemostats as opposed to flasks. In addition, we show that biofilms may be a consistent source of migrants into planktonic populations. We also show that secondary biofilms rapidly develop, although these may be more susceptible to environmental conditions. Taken together, our data suggest that chemostats may be a flexible and insightful platform for the study of biofilms *in vitro*.

**IMPORTANCE:** Biofilms are the predominant way that bacteria live in natural environments and are characterized by three emergent properties: ubiquity, resilience, and impact. They can be found across all environments, both natural and human-made, and across all of recorded time, dating back at least 3.5 billion years. Biofilms also represent a major economic impact of over $5 trillion annually. Yet, most of what is known about bacteria is the result of studies using planktonically growing liquid cultures under laboratory conditions. Here, we propose a comprehensive experimental platform that allows for study of biofilms on the molecular, organismal, and community levels using chemostats.

## INTRODUCTION

Biofilms are collections of bacteria that grow together in a dense colony, usually attached to a surface (1). They can contain one or more interacting species and are now recognized as *the* dominant way that bacteria grow in nature (1–3). Laboratory studies of planktonic growth have been incredibly successful, but they falter when we seek to better understand how bacterial communities live in natural, i.e., non-laboratory, settings.

The reasons for studying biofilms extend beyond attempts to query microbes under more naturally occurring conditions. By living in such dense and compact communities, biofilm-associated bacteria by necessity must interact, compete, and cooperate (4). These interactions create nuances that complicate our ability to fully understand microbial population biology.

Living as biofilms also allows for emergent characteristics that alter the viability of the biofilms themselves. Primary among these emergent characteristics is biofilm resilience.

Biofilms are phenomenally resilient, likely more so than any other life form in biological history. The oldest known fossils, dating back more than 3.5 billion years, are fossils of biofilm-associated communities (5). Growing as a biofilm community provides bacteria with significant protection from the outside environment, including shelter against (and even immunity from) killers such as toxins, antibiotics, viruses, and even predators.

Interestingly, the resilience of biofilms is a consequence of the makeup of these communities themselves. Being in the biofilm protects individual organisms from the outside environment while also creating a beneficial, albeit limited, internal environment. This internal environment is much like a medieval city, walled off from the outside and complete with markets for trading resources. This allows the community to survive and grow even in the face of intense external adversity, leading to extreme resiliency. Resilience is such a dominant emergent phenotype that it allows biofilms to resist damage or death over extremely long periods of time, leading to the next emergent property of biofilms, their ubiquity.

Biofilms are everywhere, existing virtually everywhere that we have ever searched for them, except for the few sterile environments that have resisted bacterial colonization. In terms of medicine and agriculture, *biofilms cause over 80% of all chronic infections in plants and animals* (6–11), highlighting once again the incredible ubiquity of biofilms and the influence they exert.

Finally, biofilms are inordinately impactful, affecting just about every part of our lives. In fact, biofilms are the direct cause of a massive *$5 TRILLION* in costs *annually* across all industries, exceeding the economic impact of *every nation on Earth, save China and the US* (12, 13). As already stated, their existence affects our health, along with that of our animals and crops. But biofilms also influence soil conditions (14) and waterways (15). They grow on our ship hulls (16) and in our canals (17), they clog our pipes (18), rot our teeth (19), and stain our monuments (20), to name but a few consequences. Taken together, these three emergent properties are of import not only to biofilms themselves, but they also directly impact our efforts to manipulate, control, and eradicate them.

In recent years, several experimental models for growing biofilms *in vitro* have been developed (21, 22), many using chambered flow-cell devices (23–25), although other well-designed platforms are also in use (26). Each of these methods has specific benefits and limitations and none address the needs of every researcher or line of inquiry. This is due in no small part to multi-factorial complexities biofilms introduce to their study *in vitro*. To explore the ecological, evolutionary, and physiological properties of biofilms more deeply, we developed a chemostat-based experimental platform for studying biofilms (Fig. 1A). Chemostats have long offered a powerful method for systematically studying microbial growth and behavior to better understand the regulatory networks that control cell growth, the adaptation of microbes to altered environmental conditions, and interactions among microbial consortia, making them invaluable tools in microbial systems biology, ecology, and evolution. Here, we apply these benefits to the study of biofilms.

**Figure 1:**
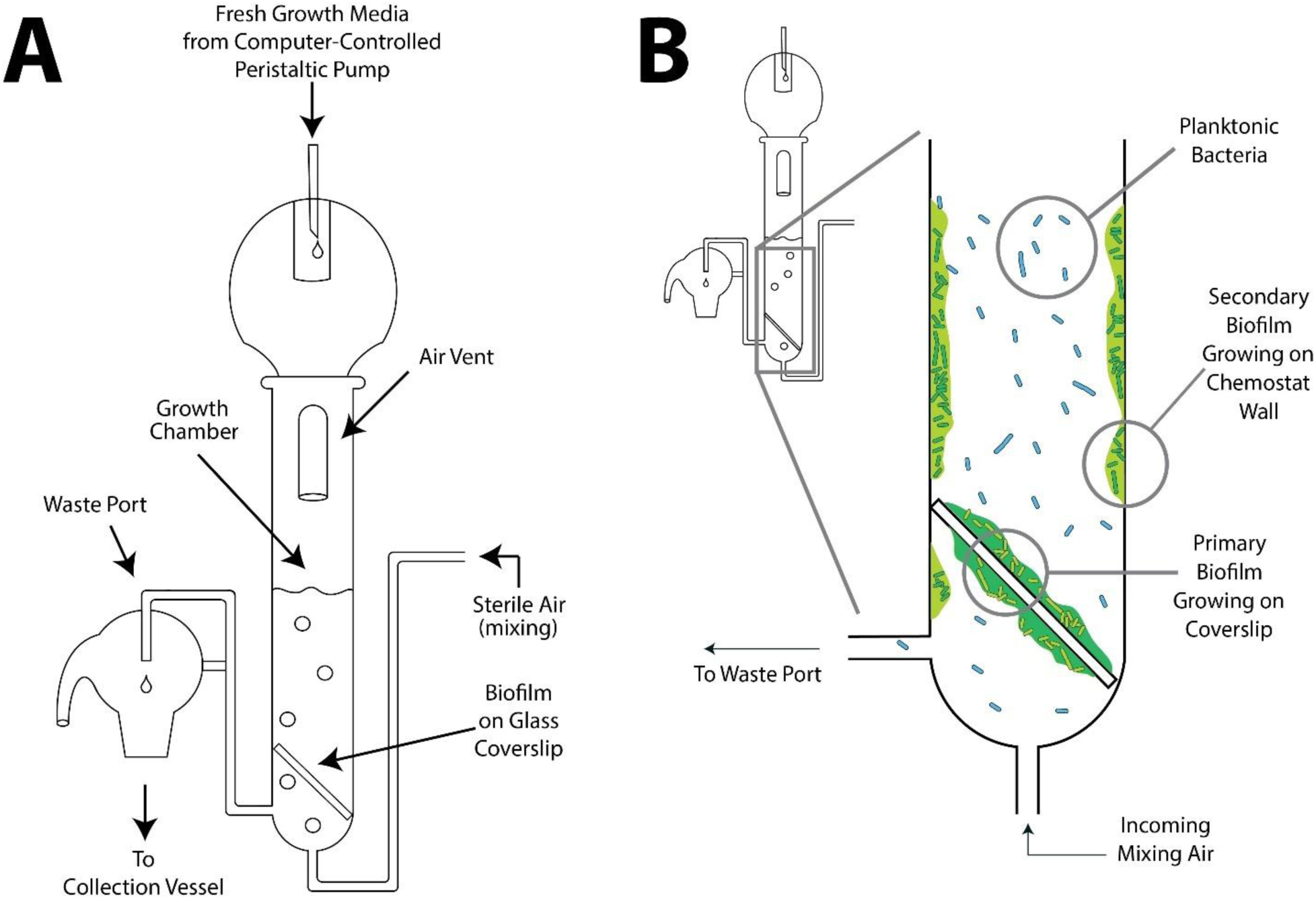
Chemostats for Biofilm Investigations. A. The biofilm chemostat, made of glass, consists of a growth chamber with a fixed volume (here, 30 mL). Fresh media is constantly added via a computer-controlled peristaltic pump and contains all nutrients needed for bacterial growth except for a single limiting concentration of a necessary nutrient. Sterile air or other gas is pumped in to mix the growth media and maintain a homogeneous liquid. A biofilm can be introduced on a glass coverslip or developed *in situ* from introduced planktonic bacteria. B. The locations of different bacterial populations that develop within the chemostat. Here, we show the primary biofilm growing on the glass coverslip, the planktonic population that develops after the seeding biofilm is introduced and the secondary biofilm that develops along the inside wall of the chemostat growth chamber.

In its simplest iteration, a chemostat consists of a closed growth chamber containing a fixed volume of growth medium, which receives an inflow of fresh media at a constant rate driven by a peristaltic pump. The inflow of fresh media is precisely matched by an outflow of spent media. Under these conditions, bacterial populations are maintained in a state of perpetual exponential growth as long as nutrients are continuously introduced. For any specific nutrient inflow rate, bacterial populations are maintained in a steady state at or near the carrying capacity of the chemostat system. Chemostat-based approaches have been used for biofilms in the past (27–29) and may offer advantages for testing specific aspects of their unique biology.

These advantages for researchers include the ability to: 1) study larger biofilm communities than is often possible in chambered flow-cells; 2) study biofilms from founding through growth and maturity; 3) use the sensitivity of chemostats to investigate metabolic shifts; 4) study both biofilm and planktonic cell populations in the same experimental context to investigate migration both into and out of the biofilm community (Fig. 1B), and 5) study newly generated secondary wall-adhered biofilm populations over time (Fig. 2). These wall-adhered biofilms have long been noted in chemostats, where they are traditionally considered problematic because they permit a heterogeneous environment within the chemostat (30, 31). Here, they are advantageous since they present an opportunity to explore another dimension of biofilm biology, namely how new biofilms are initiated from planktonic bacterial populations. However, any system designed to investigate biofilms in chemostats must provide consistency in enumerating the numerical, physiological, and ecological dynamics of biofilm-associated cells across the entire platform.

**Figure 2:**
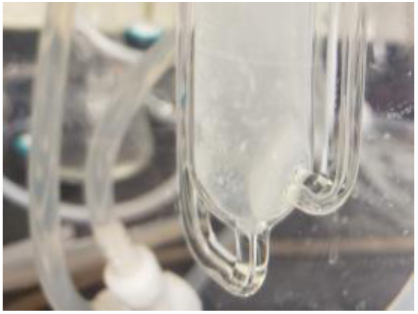
Observed Wall Biofilms. Our preliminary experiments all indicate development of large secondary biofilms on the inner walls of chemostats, obscuring the nearly clear growth media within the growth chamber. Picture taken 48 hrs after chemostat inoculation

Our method for culturing biofilms in chemostats begins by seeding chemostats with a glass coverslip on which we cultured a “nascent” biofilm. Hereafter, we refer to this nascent biofilm as the “seeding” biofilm. Once placed into the chemostat’s growth chamber, this seeding biofilm is now termed the “primary” biofilm. Bacteria emerging from the primary biofilm rapidly give rise to a planktonic bacterial population in the chemostat growth chamber as well as a secondary biofilm that forms on the walls of the chemostat. The sizes of these three sub-populations can be manipulated by altering the nutrient composition of the growth media (typically a defined medium such as Davis minimal media supplemented with glucose) or by adding biofilm growth enhancers (e.g., homoserine lactone), antibiotics, or even bacteriophages. During and after experiments, bacterial sub-population sizes can be estimated using colony counts or other imaging techniques, such as fluorescence-activated cell sorting (FACS). These techniques allow researchers to study monocultures or multi-strain biofilms, providing a powerful means of observing all phases of biofilm maturation.

We performed all experiments described here with a lab-evolved strain of *Pseudomonas aeruginosa* MPAO1, a well-studied biofilm-forming gram-negative bacterium (32) important in both ecological and clinical contexts (33). However, the chemostats are amenable to study with any number of different bacterial or fungal strains, reinforcing the flexibility of this system. Taken together, our findings suggest that chemostats can be a valuable tool to better elucidate biofilm ecology and evolution.

## MATERIALS & METHODS

The overarching goal of this work was to establish a chemostat model system as a platform for biofilm investigation. In this context, the “main experiment” describes protocols for culturing biofilms in chemostats and methods to quantify bacterial population sizes. In addition, we describe “complementary” experiments, which expand the scope of the investigations performed and highlight the versatility and applicability of this system. We also describe several “control” experiments in which we compare variations in quantification methods and begin to build best practices. The chemostat platform allows us to explore new facets of biofilm ecology, not all of which have been previously available, such as the ability to simultaneously monitor the establishment, growth, and migration of biofilm-associated cells across controlled microenvironments.

### Strains & Stocks

We performed all experiments using the lab strain, *Pseudomonas aeruginosa* MPAO1 (32). *P. aeruginosa* is a gram-negative opportunistic pathogen that is often implicated in chronic lung infections of cystic fibrosis sufferers, although it can infect several species of hosts (34) and is common across human-associated environments (35). Infections have long been recognized in both immunocompromised patients and in hospital environments (36). The PAO1 strain is a well-known *in vitro* model of *P. aeruginosa* growth and genetics (37), with MPAO1 being a sublineage with a genome size of ≈ 6.3 Mbp. In addition, both PAO1 and MPAO1 are known to be able to use hundreds of different chemical compounds as carbon sources (34) and are often resistant to multiple classes of antibiotics. Finally, *P. aeruginosa* easily forms biofilms and is a model for the study of biofilm forming organisms (33, 39).

### Pseudomonas aeruginosa FS11 Strain

*P. aeruginosa* strain MPAO1 was sourced from the Felix d’Herelle Center at the Université Laval (HER #1369). HER #1369 was adapted to the chemostat environment by growing in Davis minimal media (DMM: 1 g ammonium sulfate, 7 g dipotassium phosphate, 2 g monopotassium phosphate, 0.5 g sodium citrate, 0.2 g magnesium sulfate per 1 L water, supplemented with 0.01% glucose) for 11 days at 37°C with a dilution rate of *D*=0.28. We named this adapted *P. aeruginosa* strain “FS11” and it is the starting strain for the main experiments described here. After chemostat adaptation, we extracted FS11 DNA using the Qiagen Blood and Tissue kit and sent the purified DNA to SeqCoast Genomics (Portsmouth, NH) for whole genome Illumina sequencing. Sequencing libraries were prepared using an Illumina DNA Prep tagmentation kit and unique dual indexes. Sequencing was performed on the Illumina NextSeq2000 platform using a 300 cycle flow cell kit to produce 2x150bp paired reads. 1-2% PhiX control was spiked into the run to support optimal base calling. Read demultiplexing, read trimming, and run analytics were performed using DRAGEN v3.10.12, an on-board analysis software on the NextSeq2000. Assembly was completed using the BV-BRC platform (41). Briefly, the trimmed Illumina reads were passed through FastQ (Fastqutilities), with a Q-score threshold ≥30. Then, reads were assembled using the BV-BRC Variation Analysis software so as to be able to assemble using a GenBank genome scaffold. The Variation Analysis was run using BWA-mem-strict controls and the target genome was the reference sequence for *P. aeruginosa* MPAO1 (GenBank Accession CP027857.1; BV-BRC Genome ID 287.13733). Subsequent annotation was completed using the RAST toolkit within BV-BRC. Sequencing results were submitted to GenBank (BioSample Accession PRJNA1272633).

We prepared fresh stocks of FS11 weekly from a singular lab freezer stock maintained at -80°C in 40% glycerol. Weekly working stocks were made by streaking the frozen stock directly onto LB agar plates (10 g sodium chloride, 10 g Bacto™ tryptone, 5 g Bacto™ yeast extract, 15 g Bacto™ Agar per 1 L water) and incubating overnight at 37°C (37). We generated liquid cultures by picking a single colony from the weekly LB agar plate and inoculating 10 mL of DMM containing 0.1% of glucose as the sole carbon source (30).

### Pseudomonas aeruginosa with GFP, JJD529

A GFP-marked strain of *P. aeruginosa* was shared by the Parsek Lab at the University of Washington and was labelled as strain JJD529. This strain (Tn7::PcdrA-GFPc GmR) was modified to have a GFP tag under the control of a promoter that expresses GFP when cells are biofilm-associated. We used JJD529 for microscopy and quantitative testing in select control experiments.

### Main Experiment

#### Starting New “Seeding” Biofilms

We initiated chemostat experiments by placing a “nascent” biofilm formed on a glass coverslip, which we call the “seeding” biofilm, into the chemostat’s growth chamber. We developed seeding biofilms by placing a circular 20 mm diameter sterile glass coverslip (Cat. No. 72290-11, Electron Microscopy Sciences, Hatfield, PA) into a 50 mL conical tube containing 10 mL of fresh DMM with 0.1% glucose. A single colony of FS11 was picked from a freshly streaked LB agar plate and added to the media. The conical tube with the coverslip and bacteria was incubated overnight at 37°C shaking at 120 rpm (the conical tube was incubated at an angle of approximately 45°). The next day, we removed the coverslip, washed it in 10 mL of DMM and then transferred to 10 mL of DMM plus 0.1% glucose in a new 50 mL conical tube and incubated as described above for another 24 hours. After a total of 48 hours of inoculation, we washed the coverslip in 10 mL of sterile DMM and then aseptically transferred it to the chemostat growth chamber.

### Chemostats

All chemostats used were Kubitschek-style glass chemostats (44) with a 30 mL growth chamber volume (30). DMM with 0.01% glucose was added to the growth chamber as the only available carbon source (10x lower than in the overnight conical tubes used to start new biofilms). The addition of media into the chemostat was controlled via a Golander Model BT101F computerized peristaltic pump equipped with a DG10-4 pump head (Golander LLC, Norcross, GA). In experimental trials, the dilution rate (D) of the chemostat (30) was controlled via this pump, with trials being either “low dilution” (140.6 μL/min, D= 0.28) or “high dilution” (333.3 μL/min, D= 0.67). Temperature was maintained at 37°C using an enclosed heated water tank that housed the chemostats during all experiments, with water temperature maintained by a Lauda Type Alpha computer-controlled recirculating water thermostat (Lauda Dr R Wobser, GMBH, Lauda-Königshofen, Germany).

To ensure proper operation before experiments were initiated, we ran all chemostats for 24 hours prior to inoculation. Before adding seeded biofilms to chemostats, we drew a sample of liquid media from the chemostat and plated it on LB agar to detect any pre-inoculation contamination. To inoculate the chemostat, we removed the glass coverslip with the nascent “seeding” biofilm from the conical tube using sterile forceps, washed it in 10 mL of DMM to remove any loose cells, and then added it to the chemostat growth chamber. We drew replicate samples from the chemostat within 1 hour of inoculation and plated them to have a t_0_ measure of planktonic FS11.

We ran chemostats for 96 hours of biofilm growth. At 96 hours, final planktonic samples were drawn for quantification, the coverslip with the primary biofilm was removed and processed to determine its population size, and the secondary biofilm that grew along the inside walls of the chemostat’s growth chamber was collected and likewise quantified. We fully sequenced these populations in the same manner as previously described.

### Daily Sampling of Planktonic Cells via Serial Dilution & Plating

We drew samples from the experimental chemostats at 24-hour intervals for the 96 hours of the chemostat experiment. These samples were diluted 100,000-fold and then plated in triplicate on LB agar plates. Plating was done by placing 100 μL of the final sample dilution into 3 mL of molten soft agar (LB agar with 55% of the Bacto™ Agar used for typical agar plates) and pour plating onto the LB plate agar. After the soft agar set, a layer of top agar (Bacto™ Agar and water) was poured over the soft agar layer to aid with ease of counting colonies.

After incubating the plates overnight at 37°C, we counted and recorded CFUs the next day using a SphereFlash^®^ automated colony counter (IUL, Barcelona, Spain) and verified via manual review. We used the average of the triplicate plates for subsequent quantitative analyses.

### Sampling Primary Biofilms on the Cover Slips

At 96 hours, we removed the glass coverslip containing the primary biofilm from the chemostat using a sterile surgical clamp. We washed this coverslip in 10 mL of DMM and placed it into a sterile Potter-Elvehjem tissue homogenizer (DWK Life Sciences, Rockwood, TN) containing 1 mL of DMM. We homogenized the coverslip into a slurry over approximately 20 seconds so as to limit extraneous heat buildup. Quantification of the biofilm slurry was identical to the quantification of the planktonic samples.

### Sampling of Secondary Biofilm

After the primary biofilm was removed, we sampled and characterized the secondary biofilms that formed along the inside walls of the chemostat using a sterile silicone brush and a serological pipet. We added 5 mL of DMM to the chemostat and scrubbed the walls with the sterile silicone bottle brush, whose bristle diameter matched that of the chemostat growth chamber (≈ 20 mm). Finally, we rinsed the brush and walls with an additional 5 mL of DMM. A total volume of 10 mL of the secondary biofilm slurry was then collected, serially diluted 100,000-fold and replicate plated to quantify population size in a manner identical to the primary biofilm and planktonic populations.

## Complementary & Control Experiments

To further develop the chemostat experimental platform for biofilms, we undertook several complementary experiments to test the versatility of the system. Additionally, we conducted several control experiments to determine the consistency of quantification protocols and techniques. Additional methods and approaches used in control experiments are detailed in the Supplemental Information.

### Homoserine Lactones

We conducted a subset of chemostat experiments where a cocktail of homoserine lactones (HSL) was added to the culture media in order to experimentally manipulate cyclic di-GMP (cdGMP) production through the *LasI* and *RhlI* quorum-sensing pathways, which have been reported to assist formation and stabilization of biofilms in numerous species (45–47). Two quorum-sensing autoinducers used were N-(3-oxododecanoyl)-L-homoserine lactone (3OC12-HSL) (Sigma Aldrich, Burlington, MA) and N-butyryl-L-homoserine lactone (C4-HSL) (Sigma Aldrich, Burlington, MA).

Briefly, we prepared master stocks of these HSLs by dissolving set quantities in dimethylsulfoxide (C4-HSL: 1,000 ng/mL and 3OC12-HSL: 2,000 ng/mL). We aliquoted these stocks into 1 mL volumes and stored them at -20°C until needed. When needed for an experimental chemostat, we thawed the HSLs at room temperature and added them to the chemostat media reservoir in one of two ratios, either 4:1 (400 ng/L of C4-HSL: 100 ng/L of 3OC12-HSL). or 2:1 (400 ng/L of C4-HSL: 200 ng/L of 3OC12-HSL). We ran HSL experimental trials in chemostats at both low and high dilution rates, as above described.

### Biofilm Size Under Serial Transfer

We describe two control experiments here. The first experiment aimed to quantify the size of the “seeding” biofilm present on the glass coverslip after 48 hours of incubation. Here, we homogenized the seeding biofilm on the coverslip into a slurry using the same protocol described above, but at 48 hours of incubation under serial transfer conditions rather than after 96 hours in the chemostat.

The second control experiment compared the size of the primary biofilms grown in the chemostat with the size of the primary biofilm if it was grown solely in media that was replaced every 24 hours. The transfer of the coverslip into fresh media every 24 hours described above was extended for 144 hours, representing the total time of growth for the primary biofilm in the chemostat (48 hours to create the “seeding biofilm + 96 hours of chemostat cultivation).

### Planktonic Inoculation

In order to better understand any effect of populations being founded by biofilms versus planktonic cells, we conducted experiments where the chemostat was inoculated not with a seeding biofilm, but rather with 1 x 10^8^ planktonic cells in exponential growth phase (1 mL, OD_600_ ≈ 0.25). In these chemostats, before adding the planktonic cells, we added a sterile 20 mm glass coverslip to the chemostat growth chamber.

Prior to inoculation, we cultured planktonic cells overnight in 10 mL Davis minimal media with 0.1% glucose at 37°C, shaking at 200 rpm. The next day, we transferred 100 μL of the overnight culture to 10 mL fresh Davis minimal media with 0.1% glucose and incubated under identical conditions. When the culture OD_600_ ≈ 0.25, 1 mL of these exponentially growing planktonic cells was added to the sterile chemostat. We then ran the chemostats for 96 hrs and sampling proceeded as described for biofilm experimental trials.

## RESULTS

### Sequencing of Chemostat-Adapted FS11 Strain

Whole-genome sequencing revealed that FS11 has a single nucleotide polymorphism (SNP) in the *fliF* gene (flagellar membrane-supra-membrane ring protein), part of the *fliEFG* operon (48), causing a premature stop at codon 224. Because *fliF* is part of an operon and the MS-ring protein for which it codes is essential for the formation of the flagellum, this SNP leads to subsequent loss of the entire flagellum in FS11 (49). We verified flagellar loss using transmission electron microscopy (Fig. S1). Loss of flagella is not uncommon in adaptation to chemostat conditions, where the environment is constantly mixed and flagella provide no fitness advantage (50–52).

### Comparing Sizes of Nascent “Seeding” Biofilms with Control Biofilms on Coverslips

Seeding biofilms, i.e., those present after 48 hours of incubation in conical tubes, resulted in biofilms with an average size of 4.01 x 10^7^ CFU (95% CI: ± 2.72 x 10^7^) on the 20 mm diameter coverslips (Fig. 3, first column). Seeding biofilms were statistically smaller than the primary biofilms after 96 additional hours of cultivation in the chemostats (pairwise comparisons, Fig. 3) under all conditions except when compared to the highly variable result obtained for the high-dilution/no homoserine lactone condition, where the p-value approaches, but does not surpass the threshold, for significance (p=0.065, Wilcoxon two-sample test (53)).

**Figure 3:**
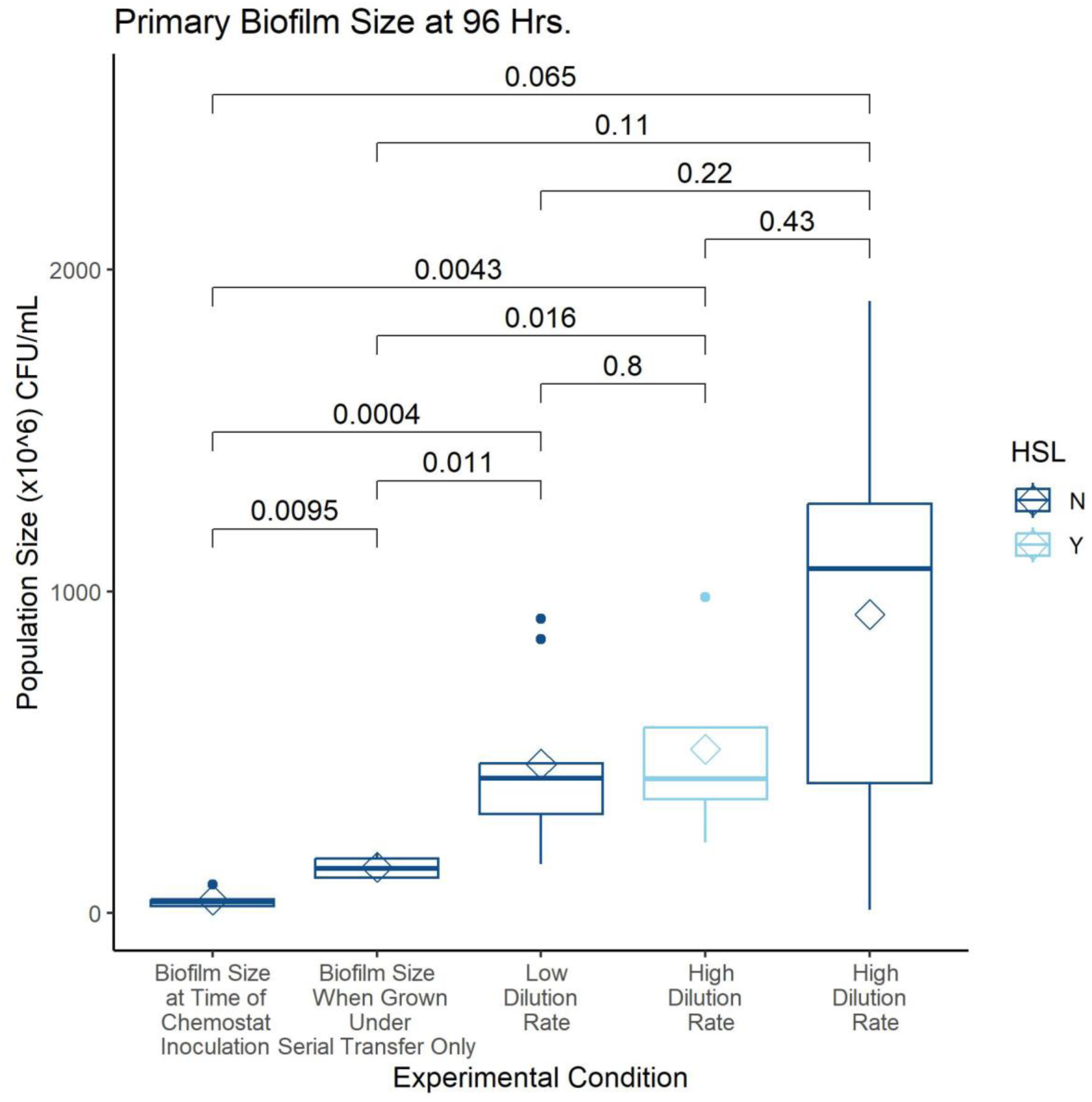
Sizes of Primary Biofilm Populations under Various Experimental Conditions. The sizes of all biofilm populations differ significantly from the size of the biofilm on the coverslip at the time of chemostat inoculation, with the exception of the high dilution/No HSL condition. In this condition, the sizes of primary biofilms were more variable, with extremely small and large sizes expanding the range of the data, yet, even in this case, the difference compared to the inoculation size biofilms approaches statistical significance (p = 0.065). The size of primary biofilms from within the chemostat are not significantly different from each other at 96 hours, regardless of chemostat environment conditions such as the dilution rate or whether HSLs are present or absent. However, all primary biofilms were larger than the seedlings or primary biofilms grown under serial transfer conditions for a total of 144 hrs. Pairwise p-values reported are the results of Wilcoxon two-sample tests following ANOVA (53).

As expected, seeding biofilms were also significantly smaller than the control biofilms grown for 144 hours under serial transfer conditions (p=0.0095, Wilcoxon two-sample test). Additionally, these control biofilms were themselves significantly smaller than those grown in chemostats, with the same exception of the high-dilution/no HSL condition (Fig. 3). These results indicate that biofilms can and do grow under serial transfer conditions but grow more steadily and to a greater extent in chemostats under all conditions evaluated.

In all experiments performed, three populations would rapidly develop. These populations were distinct but interacting and included what we call the primary biofilm on the glass coverslip, a population of planktonically growing bacteria, and a secondary biofilm population that grew on the inner walls of the chemostat’s growth chamber (Fig. 2).

### Main Experiment: Sizes of Populations Within Experiments

The size of the primary biofilm after 96 hours in the chemostat within replicate trials of a single experimental condition were consistent (Fig. 3). There was an observed increase in variation in the high dilution/no HSL condition, but this increase was not significant at the p < 0.05 level (Wilcoxon two-sample test), although all chemostat-grown primary biofilms showed high variability (Fig. 3).

Planktonic populations rapidly and consistently grew to large sizes under all conditions tested. Interestingly, all efforts to grow just a planktonic population failed as wall biofilms rapidly formed under all tested conditions. However, contrasting the size of planktonic populations to those in chemostats of other bacterial species such as *E. coli* (54, 55) indicates that the overall size of the planktonic populations of FS11 in our chemostats were comparable although such a comparison is not without inherent difficulties. For instance, these will be differences in the efficiencies of metabolism and anabolism in converting the carbon source (here, glucose) into biomass between FS11 and *E. coli* (56, 57).

The chemostat platform is perhaps the only *in vitro* system that allows researchers to simultaneously characterize primary and secondary biofilms. This capacity is vital because the formation of secondary biofilms is a hallmark of biofilms in nature. Secondary biofilms, along with planktonic populations, represent both the largest and most variable subpopulations within the chemostats, but neither the variation nor size represent problematic differences as they were all statistically consistent (Fig. 5).

### Main Experiment: Sizes of Populations Between Experiments

The sizes of primary biofilm populations across all experimental conditions either differed significantly from the size of the seeding biofilm or approached significant levels (Fig. 3, all pairwise p-values indicated). This confirms the primary objective of this series of experiments: biofilms grow under chemostat conditions. The sizes of these primary biofilms did not differ from each other across experimental conditions, suggesting that while they grow, biofilms are robust to baseline environmental conditions such as dilution rate or presence/absence of HSL signaling molecules. Whether chemostat biofilms are sensitive to additional environmental manipulation will be tested in upcoming experiments.

Planktonic population sizes across experiments were also not significantly different (Fig. 4). This result is consistent with other chemostat experiments in which planktonic populations grow to an equilibrium size (i.e., carrying capacity) and are maintained at that size throughout the experiment (30), suggesting that the dynamics of biofilm growth in chemostats aligns with chemostat theory. Strictly speaking, secondary biofilm population sizes also did not differ by condition or experiment, although our data seem inclined to suggest this is possible (Fig. 6).

**Figure 4:**
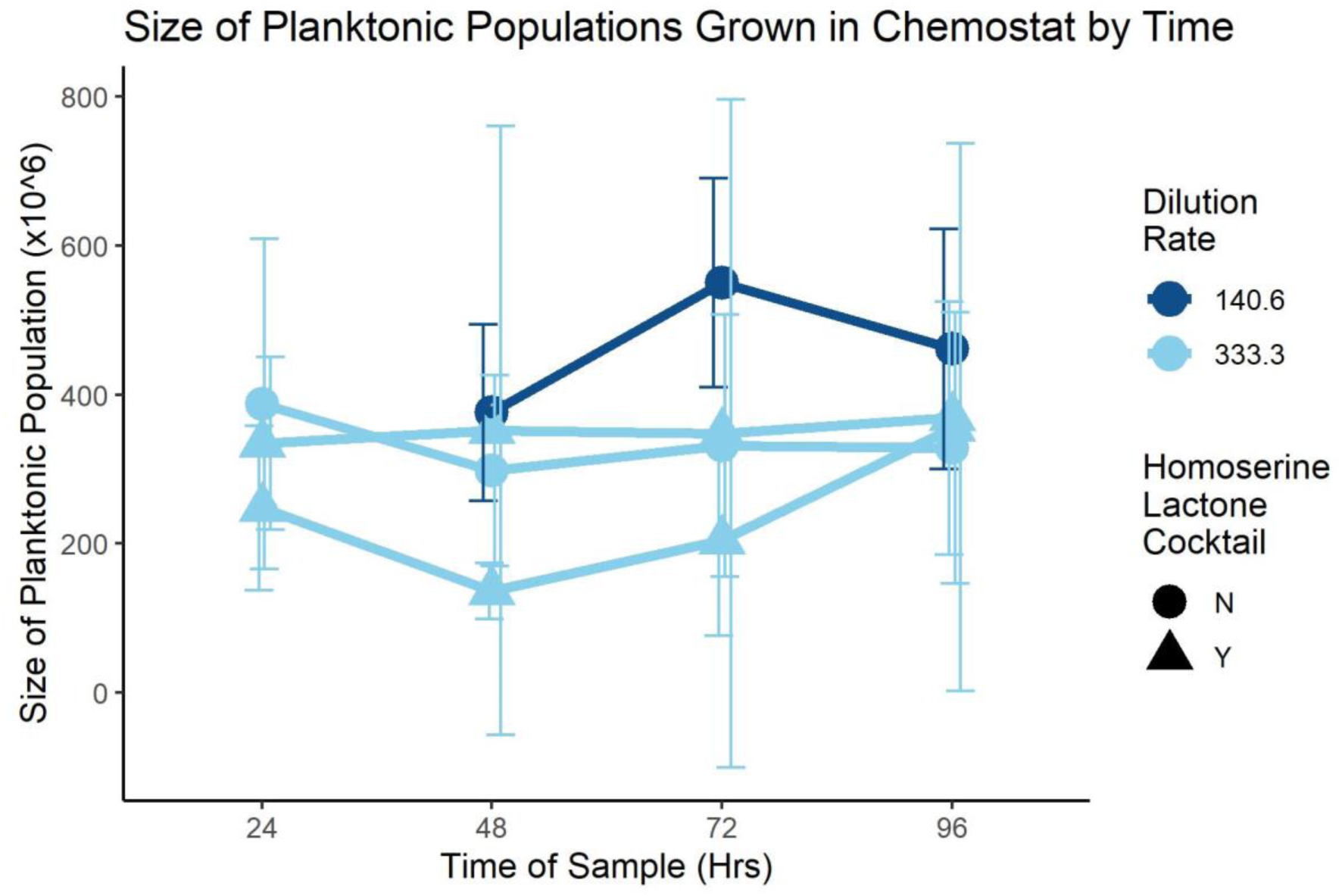
Size of Planktonic Populations in the Chemostat over Time. Several experiments were conducted and planktonic samples were taken every 24 hours after the chemostat was inoculated with the primary biofilm on the coverslip. The size of the planktonic population in CFU/mL is reported here, along with indication of whether that experiment was run with a lower rate of dilution (140.6 μL of fresh media added to the growth chamber per minute) or a high dilution rate of 333.3 μL/min. In all experiments, the planktonic population was not statistically different for each time point, suggesting that regardless of the dilution rate or the presence/absence of homoserine lactones (circles/triangles), planktonic populations were unchanged and likely under the control of the limiting nutrient, glucose. Error bars represent the 95% CI.

**Figure 5:**
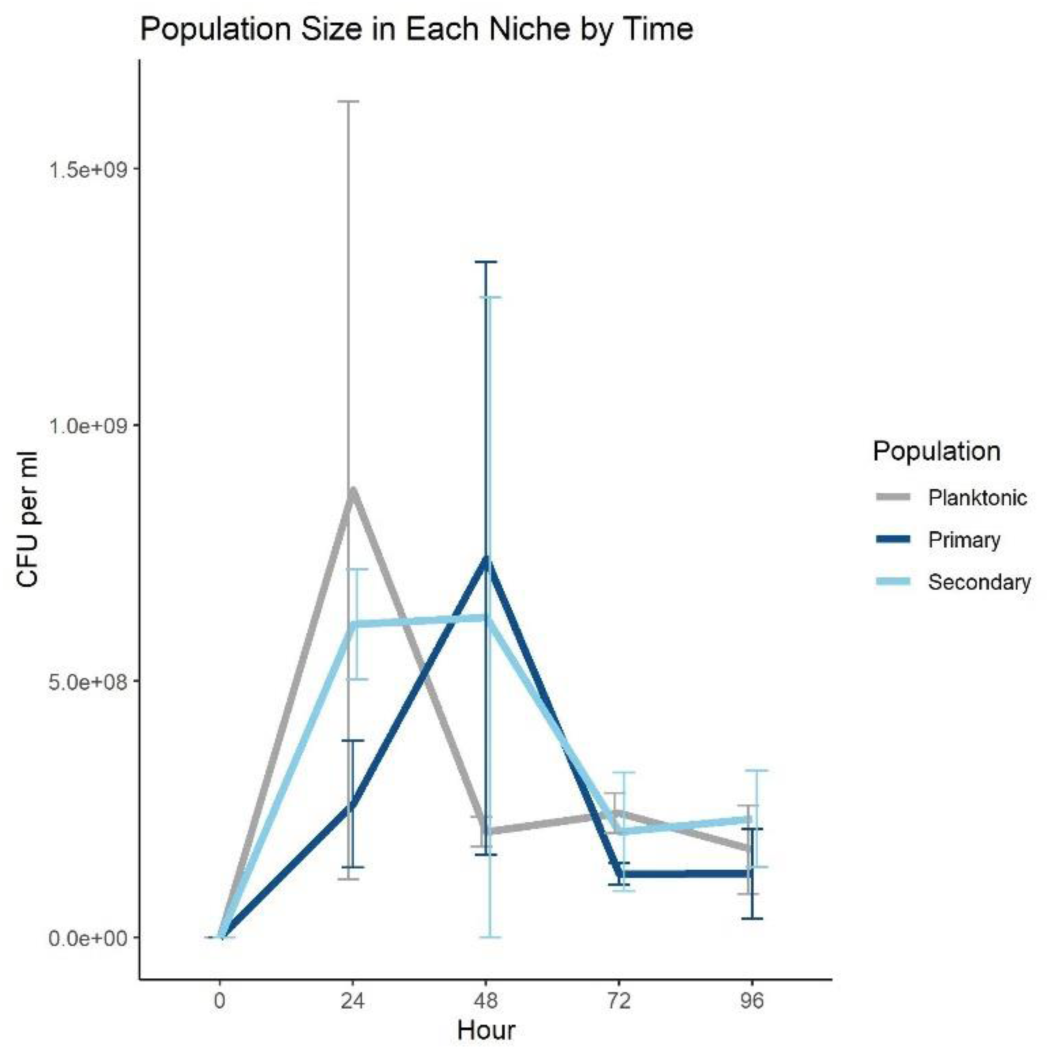
Sizes of Each Population in a Chemostat by Time. To better understand how biofilms develop within the chemostat, a control experiment as conducted in which sterile coverslips were introduced to chemostats that were then inoculated with a planktonic population of FS11. Development of planktonic, coverslip and wall biofilms were then tracked at 24-hour intervals. By 96 hours, all three populations converge to similar sizes. Note that the quantities here are in CFU/mL.

**Figure 6:**
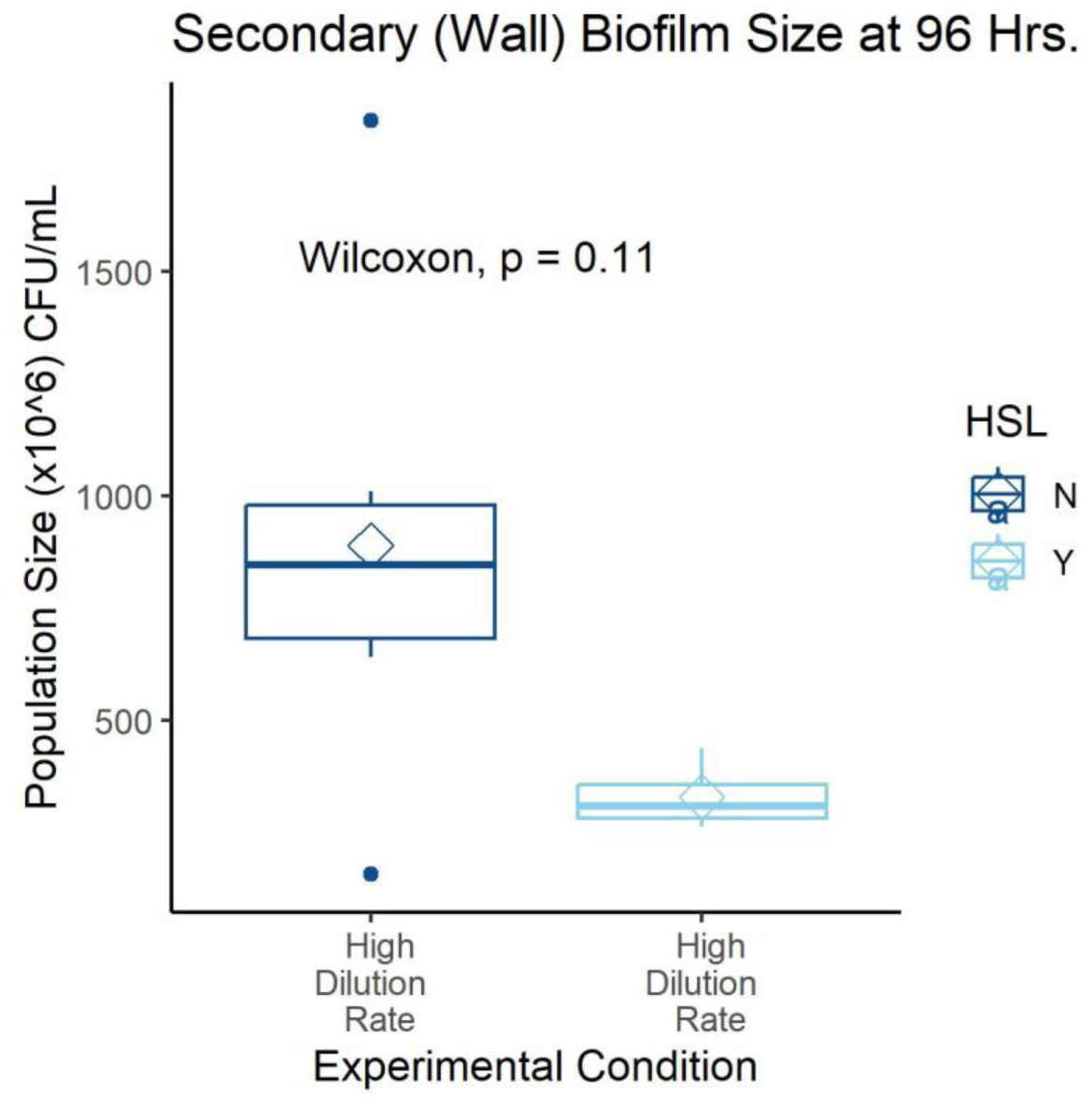
Size of Secondary Biofilm by Growth Condition. Secondary biofilms were sampled and quantified for size. While there is a qualitative difference between the HSL-conditions seen here, the data suggests that higher power may yield more informative results. One hypothesis for why this may be the case is that formation of secondary biofilm is likely to be the one most susceptible to environmental parameters because these bacteria can only come from the seeding primary biofilm and must, at least transiently, pass through the planktonic population. Once formed, a secondary biofilm can then grow, but it would be subject to the environment.

### Control Experiment: Planktonic Inoculation Experiment

In the control trials where sterile chemostats were inoculated with planktonic cells, the population sizes across all three niches (planktonic, primary, & secondary biofilms) rose rapidly, with biofilms on the coverslip lagging the planktonic and wall populations at 24 hours, only to see the sizes of the coverslip population and the planktonic population reverse at 48 hours (Fig. 5). By 96 hours, all three populations stabilized at roughly equivalent sizes. This result differs from other trials in which the chemostat is started from a primary biofilm. In those trials, the primary biofilm on the coverslip is the smallest in total population size, most likely due to the ten-fold larger area of the wall relative to the coverslip, as well as the volume of liquid media available for planktonic growth.

### Complementary Experiment: Effect of Dilution Rate and Homoserine Lactones

Several experiments were conducted to analyze the impact of media dilution rate into the chemostats. The low-dilution rate was 140.6 μL/min (D= 0.28), meaning that a volume of fresh media equal to the entire volume of the growth chamber was delivered every 3.55 hrs. The high-dilution rate was 333.3 μL/min (D= 0.67) (25), with total volume replenishment in just 1.50 hrs. No difference in the size of niche populations was observed for either final biofilm size (light vs. dark blue, Fig. 3) or in the size of planktonic populations across all timepoints sampled (Fig. 4).

In an effort to test whether chemically-signaled stabilization of biofilm phenotypes would lower the planktonic and/or secondary biofilm populations, a cocktail of autoinducer molecules involved in quorum-sensing (58, 59) was added to some experimental trials. Trials with N-(3-oxododecanoyl)-L-homoserine lactone (3OC12-HSL) and N-butyryl-L-homoserine lactone (C4-HSL) cocktails also resulted in no statistically different population sizes when compared to controls for planktonic, primary biofilm, or secondary biofilm population sizes (Figs. 3, 5, & 6).

## DISCUSSION

Biofilms have an oversized yet underappreciated impact on innumerable aspects of our modern world, from health to infrastructure to technology. This lack of recognition has left us largely uninformed as to their biology, but recently researchers have taken up the subject of how microbes live in nature and our understanding of biofilms has rapidly advanced. Here, we show that chemostats, simple culture devices that allow for continuous growth of microbial populations, can support not only biofilms, but the subsequent populations to which they give birth, making the chemostat environment closer to an “ecosystem” than to a traditional laboratory flask. Our results clearly indicate that biofilms can indeed be grown under the controlled conditions of a chemostat, that chemostats allow for multiple populations to arise, establish and be constantly supported, and, importantly, that these populations all interact. The mechanisms, mode, and manner of these interactions provide rich opportunities for further study of how biofilms grow and proliferate. In addition, we show that each of these populations can be tracked and quantified and that these approaches lead to consistent results.

Chemostats are designed to grow planktonic cells to an equilibrium size that equals the carrying capacity of the vessel and maintain that population size indefinitely (60). In this laboratory model of biofilms, planktonic populations quickly and consistently develop out of the primary biofilm. When a purely planktonic population is grown in a chemostat, the population size increases rapidly in the first hours and then levels-off to the equilibrium size. The presence of a biofilm alters this dynamic, leading the planktonic or biofilm populations to vary from trial to trial, explaining the variance in biofilm sizes observed.

Traditionally, chemostats keep (planktonic) cells growing at sub-maximal rate for indefinite periods by limiting access to a key nutrient (60). Here, that limited nutrient was glucose. The question of whether biofilms that are intentionally grown in chemostats would react in the same manner as the planktonic populations more typically grown seemed reasonable considering that biofilms are known for their resilience to harsh environmental conditions and often inadvertently grow in chemostats, causing confounding “wall effects” (30). Further, if biofilms can be grown in chemostats under controlled conditions, how would their size or constituency be influenced by populations such as planktonic cells? Lastly, given the variety of biofilm culturing methods already available to what benefit would chemostats be useful? Here, we show that chemostats provide insight into the multiple consequences of having a biofilm in the environment and begin describing some of these dynamics.

In our experiments, primary biofilms clearly grow after being inoculated into the chemostat and they are larger than the control biofilms grown via serial transfer. In fact, the difference between the sizes of biofilms grown under serial transfer and those grown in the chemostat at both low and high dilution rates suggest that serial transfer conditions are highly limiting and restrict the overall growth of biofilms. When chemostat grown, the size of the biofilm may be dependent upon the rate of carbon introduction with lower dependence upon whether or not externally applied HSLs are present (Fig. 3). The larger biofilm populations in high dilution experiments without losses in planktonic population sizes provides some insight into the resiliency of biofilms. The higher level of variation from trial to trail in the high dilution, no HSL condition for the primary biofilms also suggests that the presence of HSLs may be stabilizing.

Nonetheless, the increase in variance in the size of primary biofilms at 96 hours suggests that something biologically relevant is happening and warrants further investigation. Whether or not this phenomenon relates to shifts in community-level physiology as biofilms mature, as has long been suspected (1), remains to be determined.

In the planktonic populations, inactive or not-dividing cells would be washed out of the chemostat environment. But by their nature, biofilms retain cells within the extracellular matrix, even when they are not actively growing. In this way, biofilms become a kind of reservoir, holding higher numbers and greater genetic diversity within, even if each cell is not actively contributing to the next generation. When any environmental switch occurs, this genetic diversity may come into play, allowing the biofilm to survive and enhancing resilience. This intriguing possible hypothesis for biofilm resilience is one our platform may be particularly well-suited to explore.

The planktonic population is also as robust to changes in media dilution rate or HSL-presence (Fig. 4) as are primary biofilms, but perhaps not for the same reasons. In the high-dilution rate experiments, the average residence time of any planktonic cell in the chemostat is expected to be 1.5 hours. At such a dilution rate, chemostat theory predicts a real generation time of 1.04 hours for a purely planktonic population. However, given the additional biomass load of the coverslip and wall biofilms observed in the planktonic control experiment, we can reasonably suspect that the doubling time of planktonic cells is longer by the time the chemostat reaches the 24-hour mark. This observation would imply that a planktonic cell would be as likely to be washed out of the growth chamber as to reproduce, making it more difficult for the planktonic population to be sustained. This may explain the rapid shrinking of the planktonic population after 24 hours seen in Figure 5. This also raises the real possibility of the planktonic population being consistently replenished with cells emigrating out of the biofilms. Such a hypothesis based upon chemostats has been suggested before (25) and is the basis of concern over wall effect (30). If the biofilms are replacing planktonic cells lost, the implications may be substantial for understanding biofilms in many respects, from dispersion to robustness.

Our data also suggest that secondary biofilms may be more vulnerable to environmental influences, such as HSLs, than primary biofilms. The wall biofilms show a difference in population size when HSLs are externally applied versus absent. While the limited number of trials conducted here do not reach the level of statistical significance (p=0.11, Wilcoxon two-sample test), they do trend in that direction and the difference is evident on a qualitative level (Fig. 6). This suggests that the overall impact of the environment may be larger for the founding of subsequent biofilms than for an original biofilm itself. How biofilm-associated cells go one to found new biofilms has important implications across a range of disciplines (18, 61, 62).

Although this will not be definitively understood without significant further investigation, the possibility of using this chemostat platform regarding biofilm ecology as well as for controlling the spread of biofilms in critical applications opens several interesting avenues of inquiry.

Likewise, the chemostat platform described here holds potential as a method for investigating the role of persistent cells in the longevity and resiliency of biofilms. Persistent cells survive antibiotic and environmental insult through spontaneous phenotypic differentiation in which they are not actively growing (63). When coupled with the physical resilience of biofilms (62), the ability of persistent cells to continue on amplifies their impacts. The opportunity to design studies in which environmental conditions can be shifted and biofilm responses observed, or to just observe biofilms over more extensive periods of time, can be of great benefit to our understanding.

Lastly, although the experiments described here all contained only a single bacterial strain, the potential for multispecies studies is clear. Most naturally occurring biofilms are multispecies; the chemostat platform introduced here allows for investigation of multispecies biofilms under a range of environmental conditions, allowing researchers to go beyond the confines imposed by Koch and study bacteria *as they are in nature*. This also suggests a potential role for chemostats in the study of both multispecies and synthetic communities, particularly in relation to cross-feeding (64), or community assembly (65, 66), or other aspects of biofilms relevant to biomedicine (67), technology, and infrastructure.

Much work remains before chemostats become a dominant platform for biofilm research. The approaches introduced here also allow us to study long-neglected (and often looked-down upon) populations such as those that grow on the walls of chemostats in a quantitative and, eventually, model-ready way. However, the baseline experiments reported here provide not only a foundation for further study, but a basis for the development of entirely new approaches for the study of microbial populations and communities.

## ACKNOWLEDGEMENTS

This work was supported by the NIH (Award # 5R21AI156798) and the PSC-CUNY (Award # 66684-00 54) to JJD and start-up funds to FS. We thank George O’Toole for helpful discussions. We thank Matthew Parsek and Xuhui Zheng for generously sharing GFP-tagged strains. Special thanks to Dan Dykhuizen for comments on an earlier version of the manuscript.

